# misoTar: A novel approach for predicting miRNA and isomiR targets

**DOI:** 10.64898/2026.05.08.723919

**Authors:** Rony Chowdhury Ripan, Xiaoman Li, Haiyan Hu

## Abstract

Understanding the interactions between microRNAs/isomiRs and mRNAs has long been a major challenge in RNA biology. Although numerous computational approaches have been developed to predict these interactions, most fail to account for isomiR mediated targeting. To address this limitation, we developed misoTar, a deep learning framework trained on more than 6.662 million positive and negative interaction pairs derived from 67 publicly available human samples across six independent studies. In five-fold cross-validation, misoTar achieved an average precision of 0.930 and a recall of 0.898. Evaluation on independent test datasets demonstrated consistently superior or comparable performance relative to existing tools, including TargetScan, Mimosa, DMISO, and TEC-miTarget. In addition, single-nucleotide mutation analyses of true positive interactions revealed the critical functional contributions of non-seed regions in microRNA/isomiR targeting. Overall, misoTar provides a robust and accurate framework for predicting microRNA/isomiR interactions while offering new biological insights into microRNA targeting mechanisms. The misoTar tool is publicly available at https://figshare.com/projects/misoTar/262723.

## Introduction

MicroRNAs (miRNAs) are ∼22 nucleotide (nt) non-coding RNAs that play essential roles in post-transcriptional gene regulation^1^. They typically bind to 3′ untranslated regions of target mRNAs, leading to translational repression or mRNA degradation^2^. This interaction has traditionally been attributed to the conserved seed region (positions 2–8)^3^, although recent evidence shows that non-canonical interactions involving mismatches, bulges, and 3′ compensatory pairing can also be functional^4–6^. Dysregulation of miRNA–mRNA binding has been implicated in numerous diseases^1,7–10^, underscoring the importance of accurately characterizing miRNA–mRNA interactions^11^.

IsomiRs are miRNA isoforms generated during miRNA biogenesis through imprecise cleavage of pri- or pre-miRNAs or via nt additions, deletions, and substitutions^12^. These modifications produce sequence variants that differ in length and composition from their reference miRNAs^13^. Like canonical miRNAs, isomiRs bind to mRNAs to mediate translational repression or mRNA degradation^14^. However, variations in sequences, particularly in seed regions, can result in distinct target profiles^15^. This highlights the need to investigate isomiR–mRNA interactions to fully understand the regulatory complexity of miRNA biology

A wide range of computational approaches has been developed to predict miRNA–mRNA interactions^16–22^. Early methods such as TargetScan^17^ and miRanda^19^ focused primarily on conserved seed complementarity, which effectively identifies canonical sites but often fails to capture non-canonical ones. To address this issue, non-conservation-driven models like TarPmiR^16^, RFMirTarget^23^, miSVR^19^, and miRDB^24^ incorporated non-conserved sites and other features into machine learning frameworks using static representations and algorithms such as random forests and support vector regression. The emergence of deep learning further advanced the field by enabling automatic extraction of informative patterns directly from raw sequences. Models such as DeepMirTar^25^, miRAW^26^, miTAR^27^, GraphTar^28^, deepTarget^29^, and TargetNet^30^ significantly improved predictive performance by learning complex interaction patterns directly from sequence data. More recently, transformer-based methods like Mimosa^18^, miTDS^31^, and TEC-miTarget^32^ have demonstrated state-of-the-art accuracy by effectively modeling long-range dependencies and base-pairing patterns, without relying solely on seed-based heuristics. However, these tools do not differentiate between miRNA–mRNA and isomiR–mRNA interactions, which may limit their predictive accuracy. To address this gap, DMISO^33^ combined convolutional neural networks with bidirectional long short-term memory layers and was trained on miRNA-mRNA and isomiR-mRNA interactions extracted from the CLASH (Crosslinking, Ligation, and Sequencing of Hybrids) dataset^34^. Although DMISO represented an important step forward, its training on only six CLASH samples restricts its robustness and generalizability.

In this study, we extended the DMISO study and fine-tuned a Bidirectional Encoder Representations from Transformers (BERT)^35^ model on miRNA-mRNA and isomiR-mRNA interactions derived from 67 publicly available CLASH-like human samples across six studies. We evaluated this model, which was termed as misoTar, using five-fold cross-validation and independent datasets. misoTar achieved an average accuracy of 0.915, a F1 score of 0.914, and an Area Under the Receiver Operating Characteristic Curve (AUROC) of 0.965. When compared with TargetScan, Mimosa, TEC-miTarget, and DMISO, misoTar demonstrated superior or comparable performance across all evaluated datasets, highlighting its strong predictive capability. misoTar will be a useful addition to the existing methods and tools for miRNA studies^36^.

## Material and Methods

### The Benchmark Datasets

In a previous study^12^, we compiled all miRNA–mRNA and isomiR–mRNA interactions from 67 publicly available CLASH-like human samples spanning six datasets: CLASH^34^, CLEAR-CLIP^37^, qCLASH^38^, eCLIP^39^, Kozar et al.^40^, and Fields et al.^41^. The DMISO pipeline was utilized to process chimeric reads in the data and extract miRNA/isomiR-mRNA interactions in each sample. After extraction, we extended the 3′ end of the mRNA segment represented in each chimeric read by 25 nts to obtain a more complete target site for the corresponding miRNA or isomiR. Interactions with extended mRNA target sites shorter than 30 nts were discarded to be consistent with previous practices^34^. The resulting extended pairs were treated as positive interactions, and duplicate interactions across datasets were removed. For each positive interaction, we generated a corresponding negative interaction by matching the same miRNA or isomiR to a different site within the 3′ UTR of the same mRNA transcript. Negative sites were required to be at least 10 nts away from the corresponding positive site and to exhibit a minimum free folding energy, as computed by RNACoFold, of less than –10 kcal/mol^42^, following previously established criteria^16,43^.

The final dataset consisted of 6,661,762 positive interactions and an equal number of negative interactions. We randomly assigned 80% of this dataset to training and used it for five-fold cross-validation of misoTar. The remaining 20% was held out as an independent test set. Additionally, we performed dataset-wise evaluation, training misoTar on five datasets and testing it on the remaining one.

Furthermore, we assessed misoTar using 10 miRAW^26^ test datasets. Each dataset comprises 548 experimentally validated positive interactions and an equal number of negative interactions at the gene-level. Because the full-length mRNA sequences in the miRAW datasets are considerably longer than actual miRNA/isomiR target sites, we standardized all sequences to a 65-nt representation, matching the average mRNA segment length used for training misoTar. To extract the 65-nt subsequences, we first aligned each miRNA to its full-length mRNA using the Smith– Waterman algorithm^18^. The 65-nt window with the best alignment score is selected as the miRNA target site. These 65-nt sequences were used as input for misoTar and DMISO, as both models consider mRNA segments for potential miRNA/isomiR target sites. In contrast, Mimosa and TEC-miTarget include their own candidate target site selection mechanisms and were therefore provided with the complete mRNA sequences. TargetScan was also evaluated using the full-length mRNA sequences available in the miRAW dataset.

We also collected experimentally validated miRNA target sites from miRTarBase (release 10.0)^44^. We removed interactions lacking corresponding miRNA and mRNA identifiers in miRBase v22.1^45^ and GENCODE v46^46^, respectively. To ensure comprehensive target site coverage, we extended the 3′ end of each mRNA sequence by 25 nts and excluded interactions whose extended sequences were shorter than 30 nts. We also excluded any interactions present in the training sets of misoTar, Mimosa, DMISO and TEC-miTarget. The final miRTarBase set contained 135,744 positive miRNA–mRNA interactions. Similarly, we collected 1,308 experimentally validated positive miRNA–mRNA interactions from miRecords^47^, which also contained no negative interactions.

#### Input Representation of misoTar

The input to misoTar consists of a miRNA/isomiR sequence and an mRNA sequence concatenated into a single token sequence. Tokenization is performed using a pretrained BERT tokenizer based on the WordPiece algorithm^48^. WordPiece tokenization segments nt sequences into variable-length subsequences using a vocabulary of frequent sequence fragments and maps each subsequence to a unique numerical identifier. In addition, some special tokens, such as classification token [CLS], and separator token [SEP], are added in the token sequence.

Each input sequence begins with the ([CLS]) token. During training, the [CLS] token attends to all other tokens in the sequence through the self-attention mechanism, enabling it to progressively accumulate contextual information from the entire sequence. The hidden state of [CLS] in the final transformer layer therefore serves as an aggregate sequence representation, which is utilized for predicting miRNA/isomiR–mRNA interactions (Fig. 1).

**Fig 1.**
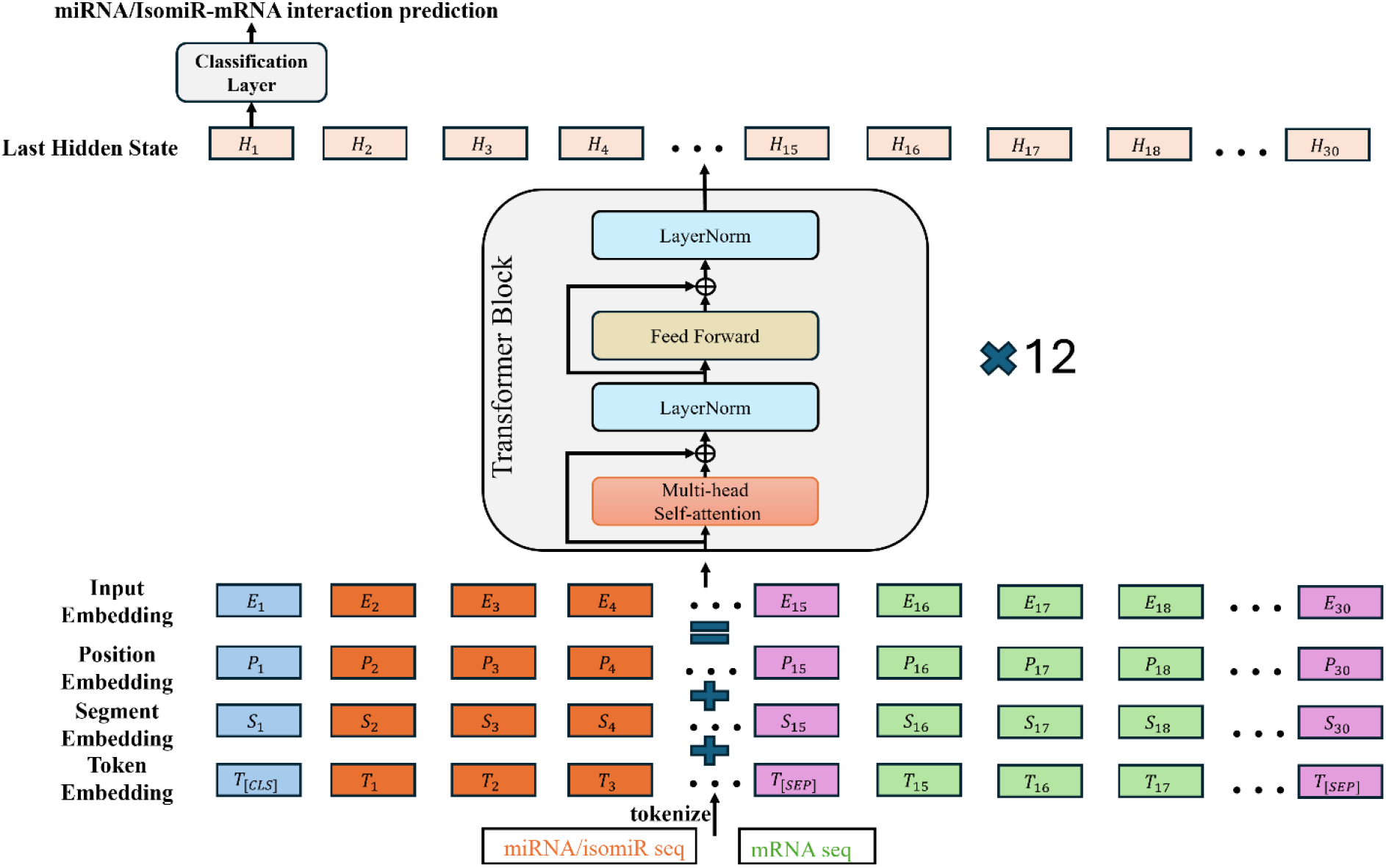
Overview of the misoTar model.

For each token, three types of embeddings are constructed: Token embeddings, Segment embeddings, and Positional embeddings. Token embeddings encode the identity of each token based on the WordPiece vocabulary. misoTar distinguishes between the two input components (miRNA/isomiR and mRNA) using both the [SEP] token and segment embeddings. The [SEP] token marks the boundary between the two sequences, while segment embeddings explicitly indicate whether a token originates from the miRNA/isomiR or the mRNA. Positional embeddings encode the position of each token within the sequence, providing information about token order, which is essential for capturing biologically meaningful sequential patterns. These embeddings are summed element-wise to form the final input embedding E (Fig. 1).

#### The misoTar model

misoTar is a BERT model that is finetuned to learn interactions between miRNA/isomiR and mRNA sequence pairs. It simultaneously captures dependencies in both left-to-right and right-to-left directions, making it well-suited for understanding relationships between paired miRNA/isomiR and mRNA.

misoTar takes the final input embedding matrix E as input, which is then passed through the encoder, consisting of a stack of identical transformer blocks. Each block includes two main sub-layers: a multihead self-attention layer and a feed-forward layer. The multi-head self-attention sub-layer is critical for modeling complex interdependencies between tokens, particularly the biologically meaningful interactions between nt positions in miRNA/isomiR and mRNA sequences. Multihead self-attention on E is computed as follows:

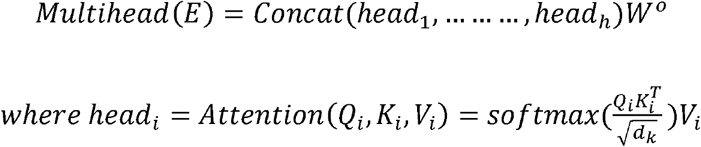

Here *Q* = *EW*^*Q*^, *K* =*EW*^*K*^ and *V* =*EW*^*V*^ are referred to as Query, Key, and Value, respectively. *Wo, W*^*Q*^, *W*^*K*^, *W*^*V*^are learnable parameters and *dk* is the dimensionality of key vectors. The feed-forward sub-layer consists of two linear layers with the rectified linear unit (ReLU) activation function, which is defined as follows:

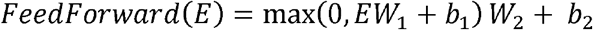

A combination of residual connections and layer normalization is applied around the multi-head self-attention and the feed-forward layer to solve the vanishing gradient problem and accelerate convergence (Fig. 1).

#### Finetuning and evaluation metrics

The BERT_BASE_ model consists of 12 transformer blocks/layers, a hidden size of 768, and 12 attention heads, totaling approximately 110 million parameters^35^. We finetuned the model using a peak learning rate of 5e-5, a batch size of 64, and the AdamW optimizer with a weight decay of 1e-2 as previously^35^. Binary cross-entropy was used as the loss function, defined as:

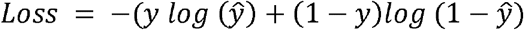

To evaluate the model’s performance, we used five commonly adopted metrics: accuracy, F1 score, precision, recall, and AUROC.

### Comparison with existing methods

We compared misoTar against four established miRNA–mRNA interaction prediction tools: TargetScan (version 7.2)^49^, DMISO^33^ and two recently developed transformer-based models, Mimosa^18^ and TEC-miTarget^32^. The evaluation was performed on the 20% held-out test dataset, ten miRAW test datasets, the miRTarBase dataset, and the miRecords dataset.

TargetScan requires separate input files for miRNA and mRNA sequences. A test interaction was considered a positive prediction by TargetScan if the miRNA ID and mRNA gene ID of the test pair matched any of the predicted interactions. In contrast, DMISO, Mimosa, and TEC-miTarget accept complete miRNA–mRNA sequence pairs as input. For these three methods, we used their default hyperparameters and the pretrained best-performing models provided by the authors. For test cases involving isomiR–mRNA pairs, we replaced the miRNA sequences with the corresponding isomiR sequences when generating the input for these models.

### Comparison with different tokenization techniques

To assess how different tokenization strategies influence model performance, we evaluated three approaches: WordPiece^48^, K-mer^50^, and Byte Pair Encoding (BPE)^51^. K-mer tokenization segments a sequence into overlapping fixed-length substrings of size k. In contrast, BPE iteratively merges the most frequent adjacent character pairs in the training corpus, producing a vocabulary of variable-length subsequences. When applied to new sequences, BPE selects the longest matching subsequences from this vocabulary and maps each to a unique identifier.

Because misoTar relies on pretrained BERT WordPiece embeddings, it cannot be directly and fairly compared to K-mer or BPE tokenization by fine-tuning alone. To ensure a fair comparison, we used DNABERT^52^ for K-mer tokenization and DNABERT2^50^ for BPE-based tokenization, as both are pretrained with their respective tokenization strategies. We selected k = 6 since DNABERT performs best with 6-mers. DNABERT and DNABERT2 were finetuned using hyperparameters similar to misoTar, a peak learning rate of 5e-5, a batch size of 64, and the AdamW optimizer with a weight decay of 1e-2.

### Feature analysis

We systematically mutated each nt position in the miRNA/isomiR sequences to determine which regions are important and crucial for sustaining miRNA/isomiR–mRNA interactions. Our analysis focused on miRNAs/isomiRs from positive interactions that were also correctly predicted as positive by misoTar (true positives). For each position, we generated four mutations: substitution with each of the three alternative nts and deletion of the nt at that position. This approach allowed us to identify important and crucial positions for each miRNA/isomiR. A position was considered crucial if none of the mutations preserved the miRNA/isomiR-mRNA interaction and important if at least one of the three types of mutations disrupted the interaction. Since a given miRNA/isomiR may participate in multiple interactions with different targets, we aggregated the important and crucial positions across all of its interactions. To avoid redundancy, positions were maintained as sets, ensuring no duplication was introduced.

## Results

### misoTar predicts miRNA/isomiR-mRNA interaction well

We performed five-fold cross-validation on the 80% training data. Among all folds, the model trained on fold 3 achieved the best overall performance during cross-validation (Table S1). We then evaluated the models trained on each fold using the remaining 20% test data. Consistent with the cross-validation results, the model trained on fold 3 also performed the best on the test set, achieving an accuracy of 0.922, an F1 score of 0.921, a precision of 0.934, a recall of 0.908, and an AUROC of 0.969 (Table 1). Across all five folds, the models exhibited strong consistency and robust predictive capability on the test data. For instance, the accuracy ranged from 0.909 to 0.922 (average 0.915), and an average F1 score was 0.914, indicating a well-balanced trade-off between precision and recall.

**Table 1.** Performance of misoTar on the 20% test data and six individual datasets.

| **Test Data** |  | **Accuracy** | **F1** | **Precision** | **Recall** | **AUROC** |
| --- | --- | --- | --- | --- | --- | --- |
| Independent<br>20% data | 1 | 0.911 | 0.909 | 0.931 | 0.887 | 0.961 |
|  | 2 | 0.914 | 0.913 | 0.928 | 0.898 | 0.964 |
|  | 3 | 0.922 | 0.921 | 0.934 | 0.908 | 0.969 |
|  | 4 | 0.909 | 0.907 | 0.923 | 0.892 | 0.961 |
|  | 5 | 0.921 | 0.919 | 0.934 | 0.906 | 0.969 |
|  | **Avg.** | 0.915 | 0.914 | 0.930 | 0.898 | 0.965 |
| Independent<br>dataset | CLASH | 0.891 | 0.887 | 0.926 | 0.851 | 0.955 |
|  | CLEAR-CLIP | 0.900 | 0.899 | 0.911 | 0.888 | 0.961 |
|  | qCLASH | 0.919 | 0.917 | 0.935 | 0.899 | 0.973 |
|  | eCLIP | 0.855 | 0.845 | 0.906 | 0.793 | 0.933 |
|  | Kozar | 0.923 | 0.920 | 0.949 | 0.892 | 0.979 |
|  | Fields | 0.679 | 0.736 | 0.624 | 0.897 | 0.782 |
|  | **Weighted Avg.** | 0.880 | 0.878 | 0.907 | 0.855 | 0.946 |

We also evaluated misoTar’s performance on each dataset, using the remaining five datasets as training data (Table 1). On the Kozar dataset, misoTar achieved its strongest performance, with an accuracy of 0.923, an F1 score of 0.920, and an AUROC of 0.979. Across the CLASH, CLEAR-CLIP, qCLASH, and eCLIP datasets, misoTar consistently maintained high performance, achieving accuracy values above 0.850. In contrast, performance on the Fields dataset was moderate, with an accuracy of 0.679. Notably, despite the lower accuracy, misoTar achieved a high recall of 0.897 on the Fields dataset.

This reduced performance on the Fields dataset may be attributed to differences in the mRNA sequence length distribution between Fields and other datasets. In fact, we compared the two groups of mRNA sequence lengths by Kolmogorov–Smirnov (KS) test^53^ and found a substantial distributional shift, with a KS statistic of 0.754 (p < 0.001). Noticed that other datasets had an average mRNA sequence length of 61 while the Fields dataset had an average mRNA length of 105.190, we normalized all datasets to a fixed sequence length of 61, truncating longer sequences and padding shorter ones with “N.” This adjustment completely eliminated the distributional shift (KS statistic = 0) and resulted in an improved prediction accuracy of 0.710. Our model thus predicted miRNA/isomiR-mRNA interactions well; however, it was the long mRNA sequences in the Fields dataset that reduced its performance.

### The misoTar model performed better than other methods

We evaluated the predictive performance of misoTar alongside four established miRNA–mRNA interaction prediction methods: TargetScan, Mimosa, TEC-miTarget, and DMISO. The assessment was conducted using the 20% held-out test dataset as well as three independent external datasets: miRAW, miRTarBase, and miRecords. For miRAW, which consists of ten separate test sets, we report the average performance in the main text and provide the results for each individual set in the supplementary material. Since miRTarBase and miRecords contain only experimentally validated positive interactions, recall was used as the sole evaluation metric for these two benchmarks.

On the 20% test dataset, misoTar showed substantially higher performance than the four comparison tools (Table 2). For instance, misoTar’s accuracy exceeded that of TargetScan, Mimosa, TEC-miTarget, and DMISO by 42.3%, 38.7%, 41.1%, and 35.0%, respectively. The much higher performance of misoTar may be largely due to the isomiR–mRNA interactions considered by the model. Among the remaining methods, DMISO achieved the second-best performance, which is consistent with its modeling of isomiR–mRNA interactions as well. Of the three approaches that do not explicitly account for isomiRs, Mimosa performed the best, likely due to its strong capability in identifying true positive interactions (Table 2). TEC-miTarget exhibited moderate accuracy but notably low recall (0.262). TargetScan showed overall moderate performance, with an accuracy of 0.499, and maintained a relatively balanced trade-off between precision and recall.

**Table 2.**
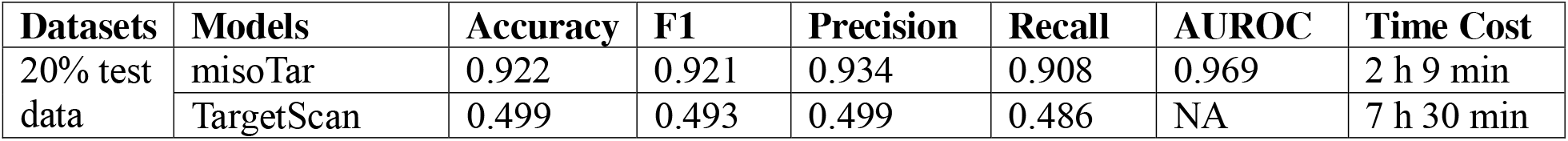
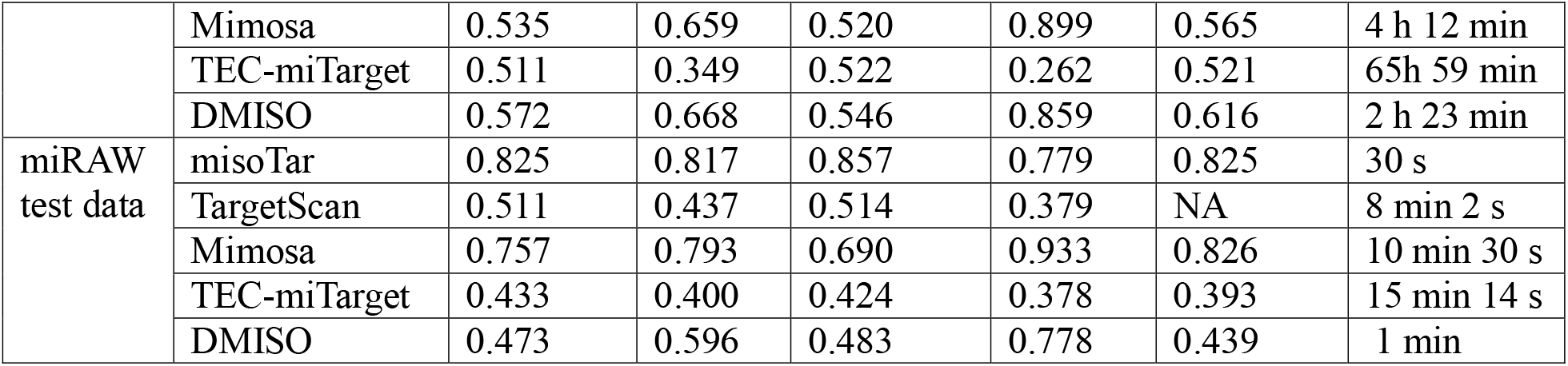
Performance comparison between misoTar with the four external tools in the 20% test data and miRAW test data.

On the miRAW test datasets, misoTar again outperformed all other tools (Table 2 and Table S2). Its accuracy exceeded that of TargetScan, Mimosa, TEC-miTarget, and DMISO by 31.4%, 6.8%, 39.2%, and 35.2%, respectively. Among the remaining models, Mimosa achieved comparatively strong performance and the highest recall (0.933). While Mimosa had a slightly higher average AUROC, misoTar showed better AUROC in five datasets than Mimosa. TargetScan, TEC-miTarget, and DMISO exhibited moderate accuracy, although DMISO achieved a notably higher recall (0.778) than both TargetScan and TEC-miTarget.

We also compared the computational cost of the models. misoTar, Mimosa, and TEC-miTarget were evaluated on a dual NVIDIA Tesla V100 16GB PCIe GPU, whereas TargetScan and DMISO were run on a 4-core Intel Core i5 CPU and a 12-core Intel Core i7 CPU, respectively. This is because TargetScan’s Pearl module and DMISO supported python 3.7.1 was unavailable on the cluster GPU we used. Notably, while misoTar, Mimosa, TEC-miTarget, and DMISO process sequences on a sample-wise basis, TargetScan performs predictions gene-wise, resulting in substantially longer runtimes. Among the deep learning models, misoTar was the fastest, followed by DMISO, Mimosa, with TEC-miTarget being the most computationally intensive (Table 2).

On the miRTarBase test dataset, misoTar achieved the highest recall, followed closely by Mimosa, with DMISO, TEC-miTarget, and TargetScan showing lower values (Table S3). On the miRecords dataset, DMISO showed a marginally higher recall than misoTar (0.774 vs. 0.771), while Mimosa performed slightly lower (0.760). Despite this small difference, misoTar maintained competitive performance and overall robustness across both datasets. Moreover, the computational cost for all models on these datasets was similar to that observed on the 20% test set, highlighting misoTar as a highly efficient and reliable predictive tool.

### The effect of parameters on the performance of misoTar

We systematically examined how various parameters affected misoTar’s performance using the 80% training data for training and the remaining 20% as a held-out test set. We investigated three tokenization strategies: WordPiece, K-mer, and BPE; as well as different model architectures and training hyperparameters.

Among tokenization strategies (Materials and Methods), DNABERT2 with BPE achieved the highest accuracy (0.928), slightly outperforming DNABERT with 6-mer (0.926) and misoTar with WordPiece (0.922) (Table S4). K-mer tokenization by DNABERT outperformed WordPiece because it better captures local sequence dependencies, which are crucial in genomic contexts. However, K-mer tokenization can introduce information leakage during masked language modeling and increases computational costs due to longer tokenized sequences^50^. In contrast, BPE tokenization by DNABERT2 functions as non-overlapping K-mers, effectively addressing this information leakage problem while creating variable-length tokens that reduce input length and enhance computational efficiency.

Despite the slightly better performance of DNABERT and DNABERT2, we chose WordPiece as the tokenization technique for misoTar. DNABERT requires 17 hours and 28 minutes to fine-tune and test, and DNABERT2 does not provide access to attention maps, limiting interpretability. Moreover, we observed unstable performance with DNABERT2 using the HuggingFace library: while folds 3 and 4 achieved high accuracy, folds 1, 2, and 5 collapsed to 0.500 (Table S5). To further investigate, we conducted five-fold cross-validations with DNABERT, which consistently produced stable results (Table S5). This finding indicates that DNABERT2 does not operate reliably within the HuggingFace implementation, likely due to incomplete or inconsistent loading of pretrained weights, leading to unstable fine-tuning outcomes.

We further evaluated misoTar with varying architectural hyperparameters, including the number of transformer layers (L), hidden size (H), and attention heads (A). Increasing model complexity consistently improved performance across all metrics (Table S6). The smallest model (L=2, H=128, A=2) showed the weakest results, whereas increasing depth and hidden size significantly enhanced predictive capability. Expanding from L=4, H=256 to L=4, H=512, A=8 yielded a notable AUROC improvement. Further increasing to L=8 and L=12 continues to improve performance, though with diminishing returns. We thus chose the model with L=12, H=768, and A=12 for a well-balanced trade-off between precision and recall.

Finally, we explored the effect of training hyperparameters, including epochs, batch size, and learning rate. misoTar achieved optimal performance with a batch size of 64, a learning rate of 5e-5, and three epochs (Table S7). Training beyond three epochs slightly reduced performance, suggesting mild overfitting. Reducing the learning rate to 2e-5 produced similar results, whereas a smaller batch size of 32 led to overfitting, as indicated by declines in precision, accuracy, and AUROC.

### The important positions in miRNAs/isomiRs

We investigated the functional relevance of individual nt positions within miRNAs/isomiRs. For each miRNA/isomiR involved in positive interactions that were correctly predicted by misoTar, we systematically mutated every position by either substituting it with each of the other three nts or deleting the nt entirely. We then assessed whether misoTar still predicted the corresponding interactions as positive. We considered a position important or crucial in a miRNA/isomiR if one or every type of mutation prevented misoTar from predicting the true positive interactions.

Our analysis revealed that important positions were most frequently located between positions 2 and 7, with a peak at position 5 (Fig. 2 and Table S8). These important positions covered beyond seed regions, suggesting the importance of both seed and non-seed regions. Crucial positions exhibited a similar but attenuated pattern. Both distributions displayed a bell-shaped trend, with a sharp decline beyond position 16. Across all indices, the number of crucial positions was consistently smaller than the number of important positions, indicating that while many positions contribute to binding stability, only a subset is essential for maintaining interaction. These results reinforce the well-known importance of the central region of miRNAs/isomiRs, particularly positions 2 to 7. Furthermore, positions observed beyond nt 24 were associated with isomiRs.

**Fig 2.**
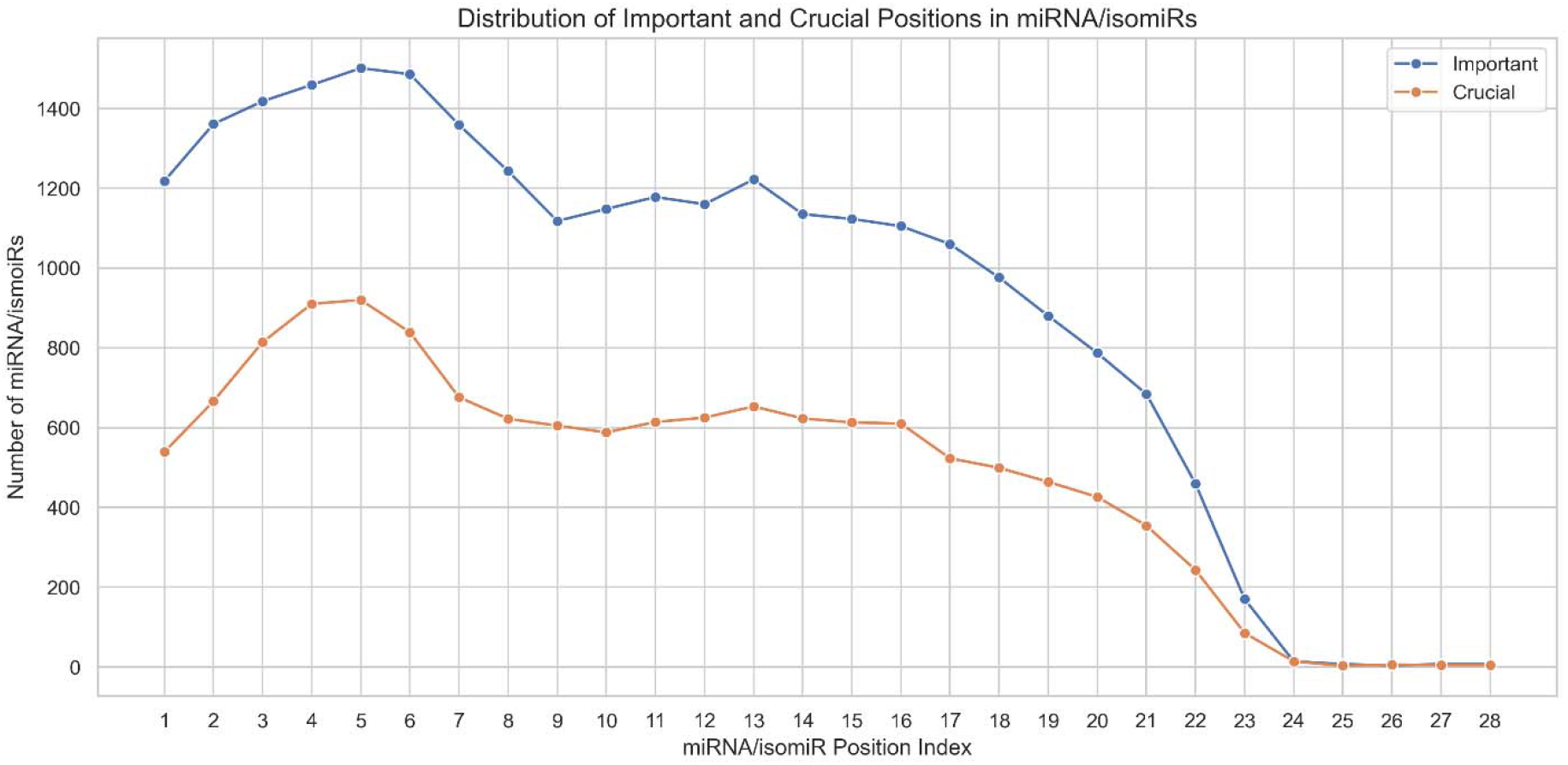
Distribution of important and crucial positions in miRNA/isomiRs.

We further classified all 2,250 miRNAs/isomiRs based on the locations of their important positions. Specifically, miRNAs/isomiRs were grouped as seed region users (all important positions within the first or last 8 nts), middle region users (all important positions between positions 8 and 15), or seed and non-seed region users (important positions spanning the seed, middle, and last 8 nts). Among the 2,250 miRNAs/isomiRs, 1,226 utilized both seed and non-seed regions, 686 relied exclusively on the seed region, 317 exclusively on the last 8 positions, and only 21 had all important positions confined to the middle region (Table S9).

We further studied the middle region users. We could only find a couple of miRNAs/isomiRs with their true positive interactions also predicted by TargetScan and stored in miRTarBase. For these miRNAs/isomiRs, we noticed that the prediction by TargetScan and the record in miRTarBase actually supported our prediction of importance positions. For instance, the isomiR of hsa-miR-19a-3p interacting with AZIN1 showed important positions at 10, 11, and 13. Although TargetScan and miRTarBase reported the hsa-miR-19a-3p and AZIN1 interaction as involving both seed and non-seed regions, miRTarBase showed pairings only at positions 10 and 13, while TargetScan showed pairing only at position 10. Therefore, the predicted important positions were supported by TargetScan and miRTarbase. The middle region miRNAs/isomiRs thus really exist and warrant future studies.

## Discussion

We developed misoTar, a tool for predicting miRNA/isomiR–mRNA interactions. misoTar reliably identified interactions with an average precision of 0.930 and a recall of 0.898. Evaluated across four independent datasets, misoTar consistently showed superior or comparable performance against existing tools, including TargetScan, Mimosa, DMISO, and TEC-miTarget. We further investigated how model parameters influenced performance and identified important and crucial positions for miRNA/isomiR binding.

Our analysis highlighted the significance of non-seed binding in miRNA/isomiR–mRNA interactions. Although more miRNAs/isomiRs have important and crucial positions within the seed region, especially position 5, the important and crucial positions span far over the seed regions (Fig. 2). Among the 2,250 miRNAs/isomiRs analyzed, 1,226 utilized both seed and non-seed regions to bind their mRNA targets. Notably, 21 miRNAs/isomiRs had important positions exclusively in the middle region (between positions 8 and 16), which is also supported by TargetScan and miRTarBase, suggesting the necessity to consider the entire miRNA/isomiR sequences for their interactions.

Although we demonstrated the good performance of misoTar, the scope of this study is limited due to the small number of available datasets and the involved miRNAs/isomiRs. With more CLASH-like samples available, we could collect more samples that provided chimeric reads of miRNA/isomiR-mRNA interactions and study more miRNAs/isomiRs. Additionally, misoTar predicts static interactions, which may not necessarily occur under specific experimental conditions or be functionally relevant. Future studies should aim to integrate experimental context and distinguish context-specific functional interactions. Finally, we documented that at least 21 miRNAs/isomiRs use only the middle regions to interact with their target mRNAs, which opens a new avenue for further study of this type of miRNA/isomiRs. Overall, our study provides a useful method and tool for miRNA/isomiR studies, which sheds new light on miRNA biology.

## Author Contributions

H.H. and X.L. conceived and designed the study. R.C.R performed the experiments. R.C.R., X.L., and H.H. analyzed the data. R.C.R., X.L., and H.H. wrote the manuscript. All authors proofread the manuscript.

## Data availability

misoTar tool and some sample data are available at https://figshare.com/projects/misoTar/262723. Our benchmark miRNA/isomiR-mRNA interaction database is available at https://figshare.com/articles/dataset/Table_S1/28958909?file=54310421. miRTarBase (release 10.0) is available at https://mirtarbase.cuhk.edu.cn/∼miRTarBase/miRTarBase_2025/php/download.php. The miRecords database used and/or analyzed during the current study may be available from the miRecords author on reasonable request. The miRAW dataset is available at https://bitbucket.org/bipous/workspace/projects/MIRAW. The CLASH dataset is available at https://www.ncbi.nlm.nih.gov/geo/query/acc.cgi?acc=GSE50452. The CLEAR-CLIP dataset is available at https://www.ncbi.nlm.nih.gov/geo/query/acc.cgi?acc=GSE73059. The qCLASH dataset is available at https://www.ncbi.nlm.nih.gov/geo/query/acc.cgi?acc=GSE101978. The eCLIP dataset is available at https://www.ncbi.nlm.nih.gov/geo/query/acc.cgi?acc=GSE198251.

